# BiGAT-Fusion: Node-Wise Gated Bidirectional Graph Attention for Drug Repurposing

**DOI:** 10.64898/2026.02.27.708641

**Authors:** Hao Chen, Chenlin Lu, Xiaoying Wei, Wenze Ding, Clara Grazian

## Abstract

Drug discovery remains slow, costly, and failure-prone, motivating computational drug repurposing that prioritizes plausible drug–disease associations (DDAs). However, DDA prediction faces three stubborn challenges: (i) extreme class imbalance with few confirmed links amid vast unknowns; (ii) directional asymmetry on the bipartite drug–disease graph that standard message passing underutilizes; and (iii) static fusion of heterogeneous evidence, where fixed rules cannot adapt the relative value of feature (similarity) and topology (association) views across nodes. We present BiGAT-Fusion, a two view graph neural model that addresses these issues end-to-end. Feature view embeddings are learned on kNN graphs built from drug–drug and disease–disease similarities, while topology view embeddings are learned via a bidirectional graph attention layer that explicitly models drug→disease and disease→drug aggregation. The views are combined by type specific, node wise gates that adaptively weight evidence per node. Pair scores are produced by a residual mixture of experts head that augments a main Multi-Layer Perceptron (MLP) with a bounded low rank bilinear residual and node biases. Under repeated *K*-fold cross validation with validation driven selection, and with the topology graph constructed strictly from training positives, BiGAT-Fusion achieves state-of-the-art AUPRC on standard benchmarks (Gdataset, Cdataset, LRSSL, Ldataset) while remaining competitive in AUROC. Analyses of learned gates and directional attentions corroborate the model’s design. BiGAT-Fusion thus offers a practical, interpretable component for large-scale, computer-aided drug repurposing.

## 1. Introduction

Drug discovery has been facing the dilemma of high investment, long period and low success rate. From design to final official approval, a new drug generally faces 90% failure probability at clinical trial phase with approximately a 15-year development cycle and 2-billion investment in advance [1, 2]. Thus, finding molecules which satisfy clinical demands while compressing the investment and cycle gradually becomes the kernel challenge in pharmaceutical field. Drug repurposing, the process of identifying novel therapeutic applications for existing medicines, has become a practical and highly effective complement to traditional de novo drug discovery [3, 4, 5, 6]. This strategy accelerates the development pipeline by leveraging compounds that already have established pharmacokinetic and safety profiles. Consequently, this paradigm can significantly reduce the attrition risk, high costs, and long development timelines typically associated with creating new medicines. In large therapeutic areas such as oncology and rare diseases alike, systematic repurposing strategies have repeatedly uncovered clinically valuable indications. For example, remdesivir is repurposed from an Ebola antiviral to a COVID-19 therapeutic [7]. However, systematic experimental testing of the enormous and largely unexplored landscape of drug–disease associations (DDA) is impractical due to the prohibitive scale involved. Powerful computational methods have therefore become essential to efficiently nominate promising candidates for experimental validation.

A central computational task in repurposing is drug–disease association prediction, where the goal is to infer plausible therapeutic links from incomplete observations. Early influential work integrated multiple similarity cues to predict novel indications (e.g., PREDICT) [8], while graph- and walk-based methods exploited heterogeneous networks of drugs and diseases, such as the bi-random-walk framework MBiRW [9]. Matrix-completion approaches, including bounded nuclear norm regularization (BNNR), further modeled the association matrix under a low-rank prior [10]. Collectively, these lines of work established that combining drug–drug and disease–disease similarity with known associations substantially improves prioritization performance. More recently, graph neural networks (GNNs) have been brought to DDA prediction to learn expressive representations end-to-end. General-purpose architectures such as graph convolutional networks (GCNs) and graph attention networks (GATs) [11, 12, 13] have inspired domain-specific designs that encode heterogeneous biomedical graphs. Representative examples include DRHGCN, which leverages heterogeneous graph convolutions [14]; DRWBNCF, which cou-ples neighborhood interaction with bilinear collaborative filtering [15]; and AdaDR, which integrates node features and topology via adaptive graph convolutions [16]. While these models achieve strong AUROC/AUPRC on standard benchmarks, there remains room to better capture directionality in bipartite message passing and to adaptively fuse complementary “feature” and “topology” views at the node level. In parallel, the quality of similarity inputs continues to matter. Drug similarity is commonly computed from 2D molecular fingerprints (e.g., ECFP) with the Tanimoto index shown to be a robust choice for fingerprint-based comparisons [17, 18]. Disease similarity is often derived from ontology- or text-based semantics (e.g., Disease Ontology/MeSH semantics) and/or functional evidence, with methods such as SemFunSim and MeSH-based semantic similarity demonstrating practical utility [19, 20]. These inputs define the “feature view” of the learning problem and complement the “topology view” constructed from observed DDAs. From a methodological perspective, DDA prediction is highly imbalanced. The vast number of unknown pairs far exceeds the confirmed positives, necessitating the use of evaluation metrics that are robust to class imbalance. In particular, under skewed class distributions, precision–recall analysis is more informative than ROC analysis, making AUPRC an essential performance measure alongside AUROC [21]. Standard evaluations typically use widely adopted benchmarks such as the Gdataset [8], Cdataset [9], LRSSL [22], and Ldataset [23].

Despite the impressive performance of recent GNN-based models, several key challenges remain to be addressed. Firstly, many existing architectures treat the drug-disease association graph with standard message-passing schemes that do not explicitly model the directional asymmetry of the bipartite interactions. The influence of a drug on a set of diseases is semantically different from the influence of a disease on the profiles of potential drugs, a nuance that is often lost. Secondly, the fusion of the feature-driven and topology-driven views is typically handled by simple concatenation or a fixed averaging scheme. Such static approaches fail to recognize that the reliability and relevance of each view can vary significantly for different nodes; for instance, a novel drug may benefit more from its feature information, while a well-studied disease may be better characterized by its rich association topology.

To address these limitations, we introduce BiGAT-Fusion, a novel two-view GNN framework designed to capture the key inductive biases of DDA prediction task. The model (i) constructs a *feature view* on intra-domain *k*NN graphs derived from drug– drug and disease–disease similarities; (ii) learns a *topology view* with direction-aware bipartite attention that explicitly models the asymmetric semantics of drug → disease and disease → drug aggregation via type-specific projections and direction-specific attentions; (iii) fuses the two views using *type-specific, nodewise gates* that adaptively weight feature versus topology evidence per node; and (iv) decodes pair scores with a *Residual-MoE* head that augments a main MLP by a bounded low-rank bilinear residual and node biases. Fig. 1 summarizes the BiGAT-Fusion pipeline, highlighting the feature/topology two-view encoders, the node-wise type-specific gates, and the Residual-MoE decoder. This design targets class imbalance, directional asymmetry on the bipartite graph, and the need for per-node fusion of heterogeneous evidence.

**Figure 1.**
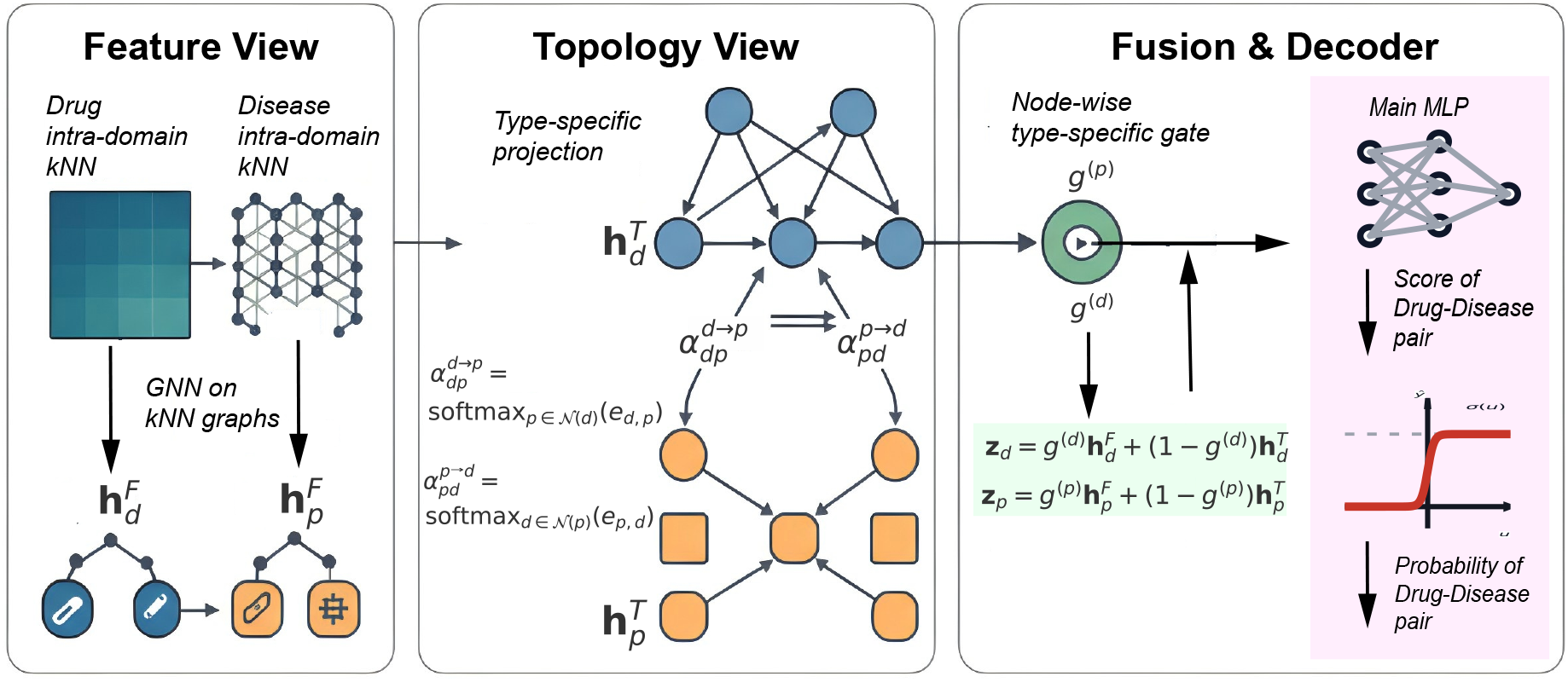
BiGAT-Fusion overview. Left: *Feature view*-GAT on intra-domain *k*NN graphs built from drug–drug and disease–disease similarities, producing 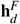 and 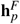. Middle: *Topology view*-bidirectional, type-specific attention on the bipartite graph with stabilized per-destination softmax, yielding 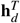 and 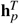 and directional weights 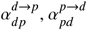. Right: *Node-wise gated fusion* with *g*^(*d*)^, *g*^(*p*)^ to obtain **z**_*d*_ and **z**_*p*_, followed by a Residual-MoE head (main MLP + bounded low-rank bilinear residual) to score drug–disease pairs.

To ensure rigorous assessment, we adopt repeated *K*-fold cross-validation with validation-driven model selection and report both AUROC and AUPRC on standard benchmarks. Our experiments show that BiGAT-Fusion achieves state-of-the-art AUPRC while remaining competitive in AUROC, highlighting the benefits of direction-aware bipartite attention and per-node gated fusion in sparse, imbalanced DDA prediction. Taken together, these contributions make BiGAT-Fusion a practically useful component for computer-aided drug repurposing pipelines: by modeling directionality on bipartite graphs and enabling per-node, data-driven fusion of heterogeneous evidence, our method addresses key bottlenecks in large-scale DDA prioritization and improves interpretability for downstream experimental validation.

## 2. Materials and Methods

### 2.1. Problem Formulation

Let 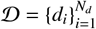 and 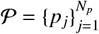 be the sets of drugs and diseases, respectively. The universe of all candidate drug–disease pairs is 𝒰 = 𝒟× 𝒫. Known associations are given by a binary matrix 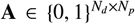, where *A*_*ij*_ = 1 indicates a therapeutic link. This induces the positive set 𝒴^+^ = {(*d*_*i*_, p_*j*_) ∈ 𝒰 | A_*ij*_ = 1} and the unknown set 𝒴^0^ = 𝒰 \ 𝒴^+^. We learn a scoring function *f*Θ 𝒟 × 𝒫 → [0, 1] such that *f*Θ(*d*_*i*_, *p*_*j*_) > *f*Θ(*d*_*u*_, *p*_*v*_) holds preferably for (*d*_*i*_, *p*_*j*_) ∈ 𝒴^+^ and (*d*_*u*_, *p*_*v*_) ∈ 𝒴^0^. For clarity, Table 1 summarizes the notation used throughout this section.

**Table 1:**
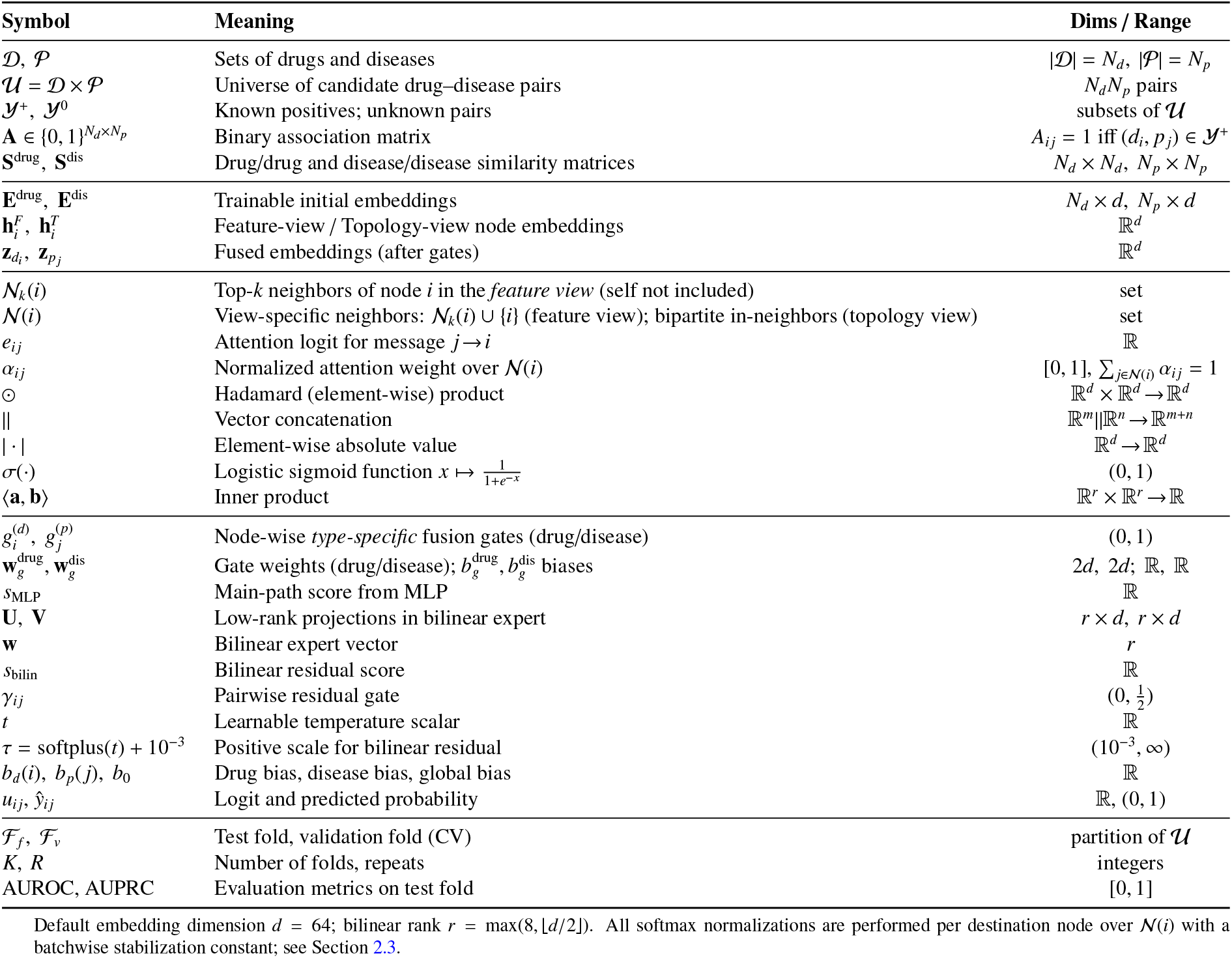
Summary of notation used in BiGAT-Fusion.

| Symbol | Meaning | Dims / Range |
| --- | --- | --- |
| $\mathcal{D}, \mathcal{P}$ | Sets of drugs and diseases | $\vert \mathcal{D}\vert = N_d, \vert \mathcal{P}\vert = N_p$ |
| $\mathcal{U} = \mathcal{D} \times \mathcal{P}$ | Universe of candidate drug–disease pairs | $N_d N_p$ pairs |
| $\mathcal{Y}^+, \mathcal{Y}^0$ | Known positives; unknown pairs | subsets of $\mathcal{U}$ |
| $\mathbf{A} \in \{0, 1\}^{N_d \times N_p}$ | Binary association matrix | $A_{ij} = 1$ iff $(d_i, p_j) \in \mathcal{Y}^+$ |
| $\mathbf{S}^{\text{drug}}, \mathbf{S}^{\text{dis}}$ | Drug/drug and disease/disease similarity matrices | $N_d \times N_d, N_p \times N_p$ |
| $\mathbf{E}^{\text{drug}}, \mathbf{E}^{\text{dis}}$ | Trainable initial embeddings | $N_d \times d, N_p \times d$ |
| $\mathbf{h}_i^F, \mathbf{h}_i^T$ | Feature-view / Topology-view node embeddings | $\mathbb{R}^d$ |
| $\mathbf{z}_{d_i}, \mathbf{z}_{p_j}$ | Fused embeddings (after gates) | $\mathbb{R}^d$ |
| $\mathcal{N}_k(i)$ | Top-$k$ neighbors of node $i$ in the *feature view* (self not included) | set |
| $\mathcal{N}(i)$ | View-specific neighbors: $\mathcal{N}_k(i) \cup \{i\}$ (feature view); bipartite in-neighbors (topology view) | set |
| $e_{ij}$ | Attention logit for message $j \rightarrow i$ | $\mathbb{R}$ |
| $\alpha_{ij}$ | Normalized attention weight over $\mathcal{N}(i)$ | $[0, 1], \sum_{j \in \mathcal{N}(i)} \alpha_{ij} = 1$ |
| $\odot$ | Hadamard (element-wise) product | $\mathbb{R}^d \times \mathbb{R}^d \rightarrow \mathbb{R}^d$ |
| $\parallel$ | Vector concatenation | $\mathbb{R}^m \parallel \mathbb{R}^n \rightarrow \mathbb{R}^{m+n}$ |
| $\vert \cdot\vert $ | Element-wise absolute value | $\mathbb{R}^d \rightarrow \mathbb{R}^d$ |
| $\sigma(\cdot)$ | Logistic sigmoid function $x \mapsto \frac{1}{1+e^{-x}}$ | $(0, 1)$ |
| $\langle \mathbf{a}, \mathbf{b} \rangle$ | Inner product | $\mathbb{R}^r \times \mathbb{R}^r \rightarrow \mathbb{R}$ |
| $g_i^{(d)}, g_j^{(p)}$ | Node-wise *type-specific* fusion gates (drug/disease) | $(0, 1)$ |
| $\mathbf{w}_g^{\text{drug}}, \mathbf{w}_g^{\text{dis}}$ | Gate weights (drug/disease); $b_g^{\text{drug}}, b_g^{\text{dis}}$ biases | $2d, 2d; \mathbb{R}, \mathbb{R}$ |
| $s_{\text{MLP}}$ | Main-path score from MLP | $\mathbb{R}$ |
| $\mathbf{U}, \mathbf{V}$ | Low-rank projections in bilinear expert | $r \times d, r \times d$ |
| $\mathbf{w}$ | Bilinear expert vector | $r$ |
| $s_{\text{bilin}}$ | Bilinear residual score | $\mathbb{R}$ |
| $\gamma_{ij}$ | Pairwise residual gate | $(0, \frac{1}{2})$ |
| $t$ | Learnable temperature scalar | $\mathbb{R}$ |
| $\tau = \text{softplus}(t) + 10^{-3}$ | Positive scale for bilinear residual | $(10^{-3}, \infty)$ |
| $b_d(i), b_p(j), b_0$ | Drug bias, disease bias, global bias | $\mathbb{R}$ |
| $u_{ij}, \hat{y}_{ij}$ | Logit and predicted probability | $\mathbb{R}, (0, 1)$ |
| $\mathcal{F}_f, \mathcal{F}_v$ | Test fold, validation fold (CV) | partition of $\mathcal{U}$ |
| $K, R$ | Number of folds, repeats | integers |
| AUROC, AUPRC | Evaluation metrics on test fold | $[0, 1]$ |

Default embedding dimension $d = 64$ ; bilinear rank $r = \max(8, \lfloor d/2 \rfloor)$ . All softmax normalizations are performed per destination node over $\mathcal{N}(i)$ with a batchwise stabilization constant; see Section 2.3.

### 2.2. Heterogeneous Graph Representation

We construct a heterogeneous graph 𝒢 = (𝒱, ℰ) with nodes 𝒱 = 𝒟 ∪ 𝒫 and two types of edges: (i) a topology bipartite graph from training positives, and (ii) feature kNN graphs inside each domain. For drugs, we build a directed *k*-NN feature graph from 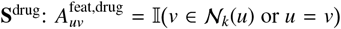 (where 𝕀 (·) denotes the indicator function), and analogously for diseases. We use *k* = 4, unless differently stated. The bipartite topology graph only uses positive training pairs 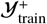 of a given fold (see Section §2.4).

### 2.3. The BiGAT-Fusion Architecture

Our model learns feature-view and topology-view embeddings in parallel and fuses them via type-specific, node-wise gates; predictions are produced by a residual mixture-of-experts (Residual–MoE) head. Concretely, we maintain trainable embedding tables for drugs and diseases, 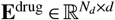 and 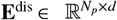, where *d* denotes the latent embedding dimension (default *d*=128). Initial node embeddings are the corresponding rows of these tables:

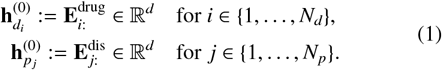

After obtaining the feature- and topology-view embeddings and fusing them into 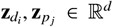, (via node-wise gates), the Residual–MoE head computes the logit for a pair (*d*_*i*_, *p*_*j*_) as

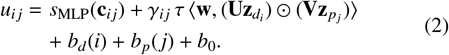

where ⊙ is the Hadamard product, ∥ denotes concatenation, τ = softplus(*t*) + 10^−3^ > 0 is a learnable temperature, 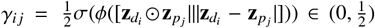 gates a low-rank bilinear residual (with projections **U, V** ∈ ℝ^r×d^ and vector **w** ∈ ℝ^r^), and *b*_*d*_(*i*), *b*_*p*_(*j*), *b*_0_ are drug, disease, and global biases. The predicted probability is ŷ_*i j*_ = σ(*u*_*i j*_).

#### 2.3.1. Layer 1: Parallel Feature and Topology Encoders Feature-View

For node *i* with neighbors *j* ∈ 𝒩_*k*_(*i*) ∪ {*i*}, attention energies are computed with the destination source concatenation,

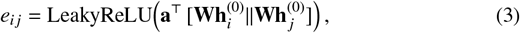

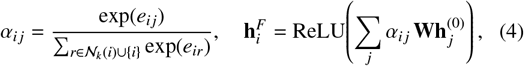

where **W** 𝒩 ℝ^*d*×*d*^ and **a** ∈ ℝ^2*d*^ are shared within each node type (one set for all drug nodes and another set for all disease nodes). Using the (destination, source) (dst, src) ordering, where the destination node is the one being updated and the source node is its neighbor providing information, aligns the implementation with the theoretical update and improves numerical stability through a per-destination softmax; we subtract a batchwise constant for stabilization (see Numerically stable normalization).

##### Numerically stable normalization

For destination node *i*, we compute attention logits *e*_*i j*_ and apply a stabilized softmax over 𝒩 (*i*) :

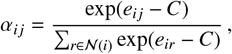

where *C* is a batchwise constant (we use the global maximum of {*e*} in the mini-batch). Because softmax is invariant to adding a constant, our formulation produces the same attention weights as the usual trick of subtracting max_*r*∈𝒩 (*i*)_ *e*_*ir*_, but it is simpler and more efficient to implement in vectorized code.

We normalize over the destination node’s incoming neighbors (per-destination softmax). Here, 𝒩 (*i*) denotes the view-specific incoming neighbor set: 𝒩_*k*_(*i*) ∪ {*i*} for the feature view, and the bipartite in-neighbors for the topology view (diseases for a drug node and drugs for a disease node).

##### Topology-View

On the bipartite graph, we pass messages in both directions with type-specific projections and direction-specific attentions:

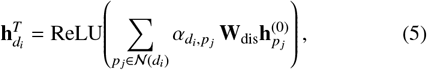

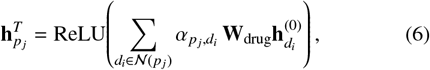

with energies

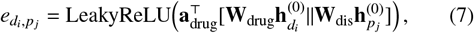

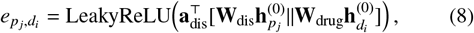

normalized per destination using the same stabilized softmax as above (see Numerically stable normalization). Here **W**_drug_, **W**_dis_ ∈ ℝ^*d*×*d*^ are type-specific, while **a**_drug_, **a**_dis_ ∈ ℝ^2*d*^ are direction-specific.

#### 2.3.2. Layer 2: Node-wise Gated Fusion

We use type-specific gates for drugs and diseases. For drug node *d*_*i*_ and disease node *p*_*j*_,

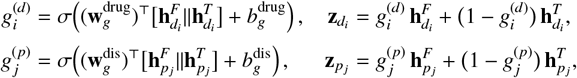

where 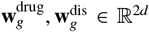 and 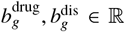. We initialize gate biases to zero, 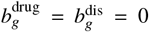, and use the PyTorch default (Kaiming-uniform) initialization for gate weights. For other projection/attention parameters we use Xavier initialization as specified in the layer definitions. This yields an unbiased starting point: under zero biases and symmetric weight initialization, the pre-sigmoid gate input has mean near zero, so σ(0) = 0.5; consequently, when the two views are on a similar scale we have 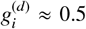 and 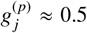for each node. By construction, 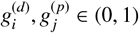.

#### 2.3.3. Residual-MoE Prediction Head

Given a pair (*d*_*i*_, *p*_*j*_) with fused embeddings 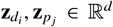, we compute the score using a main-path MLP, complemented by a bounded bilinear residual term and node-specific biases. For compactness, define the concatenations

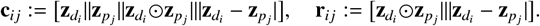

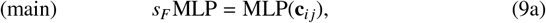

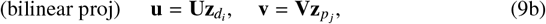

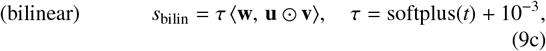

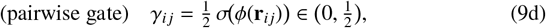

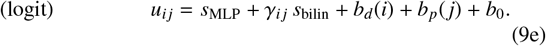

Here **U, V** ∈ ℝ^*r*×*d*^ project into a rank-*r* space (by default, *r* = max(8, *d*/2)), **w** ∈ ℝ^*r*^, *ϕ* is a small MLP, and *b*_0_ is a global bias. Constraining the gate γ_*i j*_ ∈ (0, 0.5) ensures that the bilinear expert functions as a bounded residual instead of overtaking the main path, thereby stabilizing performance as measured by AUPRC. The predicted probability is *ŷ*_*i j*_ = σ(*u*_*i j*_). We optimize the binary cross-entropy with logits over mini-batches.

Algorithm 1 summarizes the forward scoring procedure for a single pair (*d*_*i*_, *p*_*j*_) using the stabilized per-destination softmax, type-specific gates, and the Residual-MoE head described above.

##### Algorithm 1

Forward scoring for a pair (*d*_*i*_, *p*_*j*_) with BiGAT-Fusion

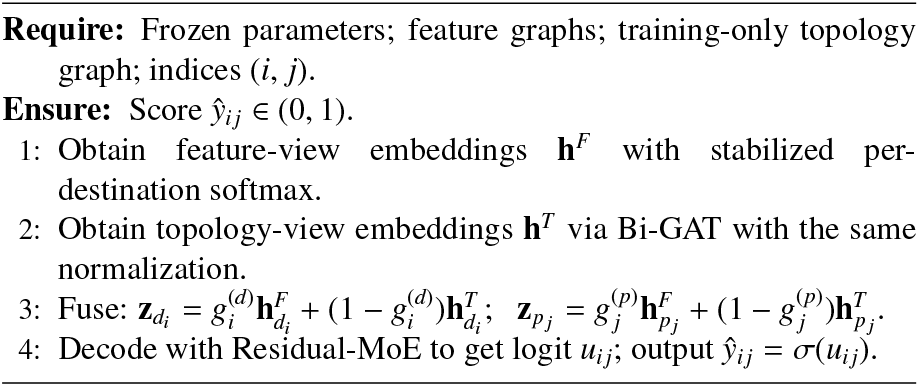

### 2.4. Training and Evaluation Protocol

#### Repeated K-fold CV

We partition the full universe 𝒰 into *K* folds 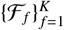 and repeat this *R* times. In each run, we hold out one fold ℱ_*f*_ for testing and another disjoint fold ℱ_*v*_ for validation. The bipartite topology graph is constructed only from positives in the remaining *K* − 2 folds (training universe), so no validation/test positives enter message passing (no structural leakage). Evaluation ranks every pair in ℱ _*f*_ (positives and unknowns) without negative sampling. On each test fold, we compute AUROC and AUPRC using continuous scores *ŷ*_*i j*_. We summarize the mean±std over 10 × 10 = 100 runs.

*Negative sampling*. In each mini-batch, *k* unknown pairs per positive are sampled uniformly from the global pool 𝒰\ 𝒴^+^. These sampled unknown pairs may originate from folds that are subsequently allocated to validation or testing. This is a standard transductive setting and does not use any label information beyond the set of known positives: no validation/test positives are used as positives during training, and no validation/test positives are inserted into the topology graph. We use a fixed negative-to-positive ratio *k*_neg_ (by default. *k*_neg_ = 3), which implicitly sets the batch class prior to π_+_ ≈ 1/(1 + *k*_neg_).

##### Algorithm 2 BiGAT-Fusion: CV training and evaluation

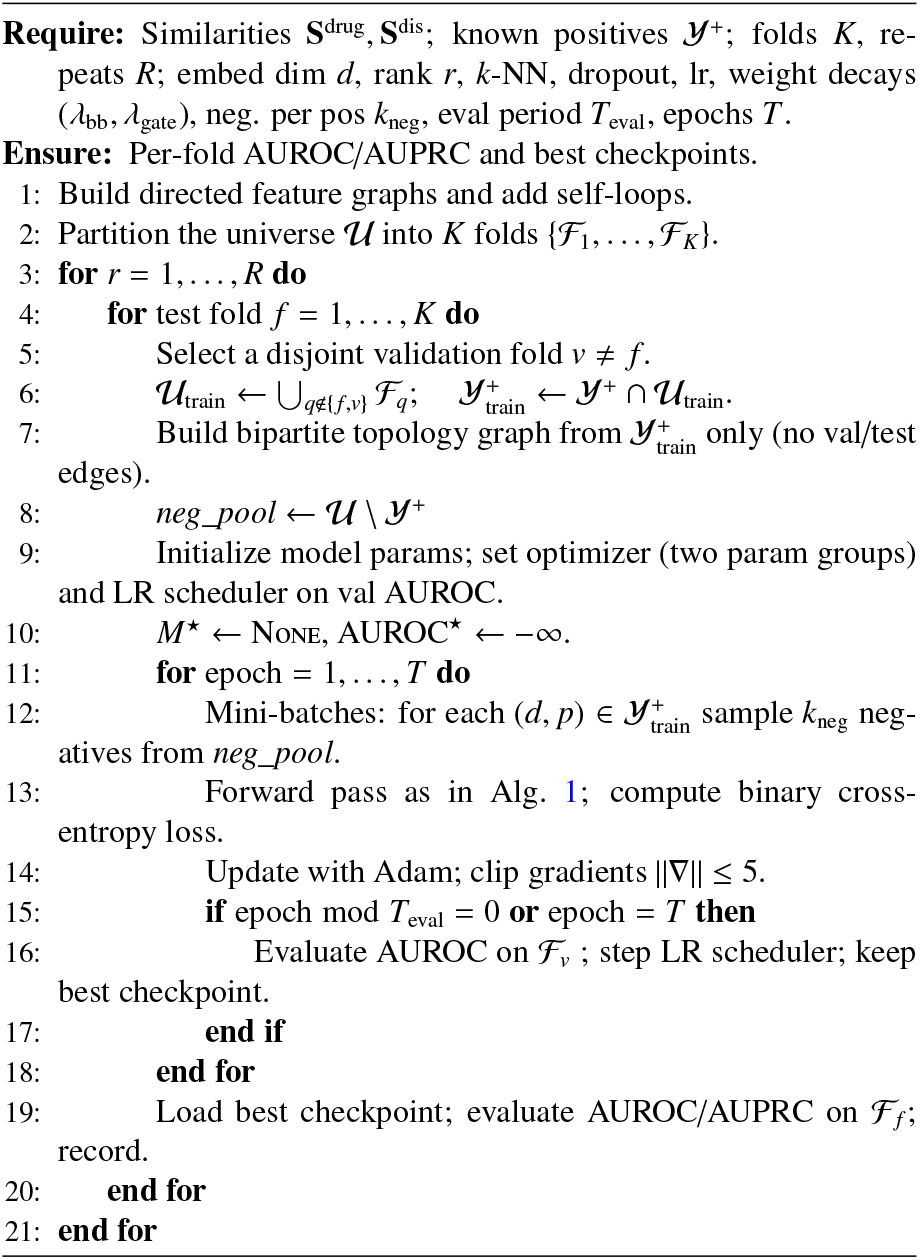

#### Mini-batch loss

We optimize the standard binary cross-entropy with logits over mini-batches, *without* per-class weighting:

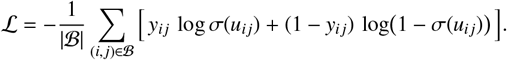

With *k*_neg_ negatives per positive, the effective class balance in a batch is controlled by *k*_neg_, which we found sufficient and stable in our experiments.

For optimization and regularization, we use Adam (lr =10^−3^ by default), gradient clipping (∥∇∥ ≤ 5), and dropout (0.2). Parameters are separated into backbone weights (all except gates), assigned weight decay 10^−4^, and gate parameters (node-fusion gates only), assigned stronger decay (10^−3^ by default) due to their calibration role. For ablation configurations (ordered by proximity to **full**), we report five controlled variants that match our implementation: (i) **full** (all components); (ii) **gate_scalar**: replace node-wise gates by a single learnable scalar per type (drug/disease); all other components identical to **full**; (iii) **topology_no_attn**: keep bipartite message passing but replace attention with a uniform mean aggregator (α_*e*_ = 1/deg(dst_*e*_)), no attention on the topology view; (iv) **feature_only**: remove the topology view; (v) **topology_only**: remove the feature view.

Algorithm 2 details our repeated *K*-fold protocol, validation-driven scheduling and testing.

## 3. Results

### 3.1. Datasets and Feature Modalities

We evaluate on four standard benchmarks: **Gdataset** [8], **Cdataset** [9], **LRSSL** [22], and **Ldataset** [23]; statistics relative to each dataset are available in Table 2. Each dataset provides (i) drug–drug similarity 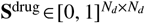 and (ii) disease– disease similarity 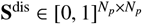. Following common practice [14, 15], drug similarity is computed from Tanimoto coefficients over 2D fingerprints, and disease similarity from MeSH term overlap.

**Table 2:** Summary statistics of the benchmark datasets.

| Dataset | Drugs $N_d$ | Diseases $N_p$ | Positives $\vert \mathcal{Y}^+\vert $ | Sparsity* |
| --- | --- | --- | --- | --- |
| Gdataset | 593 | 313 | 1,933 | 0.0104 |
| Cdataset | 663 | 409 | 2,532 | 0.0093 |
| LRSSL | 763 | 681 | 3,051 | 0.0059 |
| Ldataset | 269 | 598 | 18,416 | 0.1145 |

\* Sparsity is $|\mathcal{Y}^+|/(N_d N_p)$ .

^*^ Sparsity is |𝒴^+^|/(*N*_*d*_ *N*_*p*_).

### 3.2. Overall Performance Comparison

To rigorously evaluate our proposed model, we compare BiGAT-Fusion with seven representative baselines spanning classical and deep models: **MBiRW** [9], **iDrug** [24], **BNNR** [10], and four GNNs—**NIMCGCN** [25], **DRHGCN** [14], **DRWB-NCF** [15], and **AdaDR** [16]. All methods are re-implemented and evaluated under the same protocol of 10 repeats of 10-fold cross-validation described in Section 2.4. Aggregate results are reported in Table 3 and visualized in Fig. 2.

**Table 3:** Overall performance comparison on four benchmark datasets.

| Method | Gdataset |  | Cdataset |  | LRSSL |  | Ldataset |  |
| --- | --- | --- | --- | --- | --- | --- | --- | --- |
|  | AUROC | AUPRC | AUROC | AUPRC | AUROC | AUPRC | AUROC | AUPRC |
| MBiRW | 0.818 ± 0.021 | 0.236 ± 0.026 | 0.850 ± 0.015 | 0.364 ± 0.031 | 0.854 ± 0.015 | 0.161 ± 0.019 | 0.689 ± 0.007 | 0.256 ± 0.010 |
| iDrug | 0.727 ± 0.020 | 0.119 ± 0.022 | 0.770 ± 0.019 | 0.207 ± 0.026 | 0.717 ± 0.016 | 0.115 ± 0.017 | 0.767 ± 0.005 | 0.339 ± 0.013 |
| BNNR | 0.806 ± 0.027 | 0.490 ± 0.040 | 0.836 ± 0.021 | 0.580 ± 0.030 | 0.786 ± 0.019 | 0.389 ± 0.030 | 0.862 ± 0.004 | 0.506 ± 0.013 |
| NIMCGCN | 0.910 ± 0.014 | 0.375 ± 0.052 | 0.933 ± 0.014 | 0.459 ± 0.079 | 0.913 ± 0.011 | 0.262 ± 0.033 | 0.851 ± 0.005 | 0.508 ± 0.013 |
| DRHGCN | 0.889 ± 0.016 | 0.441 ± 0.043 | 0.909 ± 0.013 | 0.535 ± 0.056 | 0.887 ± 0.012 | 0.320 ± 0.036 | 0.872 ± 0.005 | 0.568 ± 0.018 |
| DRWBNCF | 0.908 ± 0.015 | 0.576 ± 0.039 | 0.925 ± 0.011 | 0.639 ± 0.045 | 0.898 ± 0.011 | 0.453 ± 0.030 | **0.880 ± 0.008** | 0.632 ± 0.008 |
| AdaDR | 0.895 ± 0.015 | 0.517 ± 0.046 | 0.917 ± 0.011 | 0.576 ± 0.045 | 0.888 ± 0.013 | 0.407 ± 0.054 | 0.877 ± 0.004 | 0.577 ± 0.007 |
| **BiGAT-Fusion** | **0.945 ± 0.011** | **0.647 ± 0.045** | **0.960 ± 0.008** | **0.705 ± 0.045** | **0.942 ± 0.008** | **0.516 ± 0.035** | 0.879 ± 0.004 | **0.673 ± 0.015** |

Results are reported as mean ± std. dev. The best result in each column is in **bold**, and the second-best is underlined.

**Figure 2.**
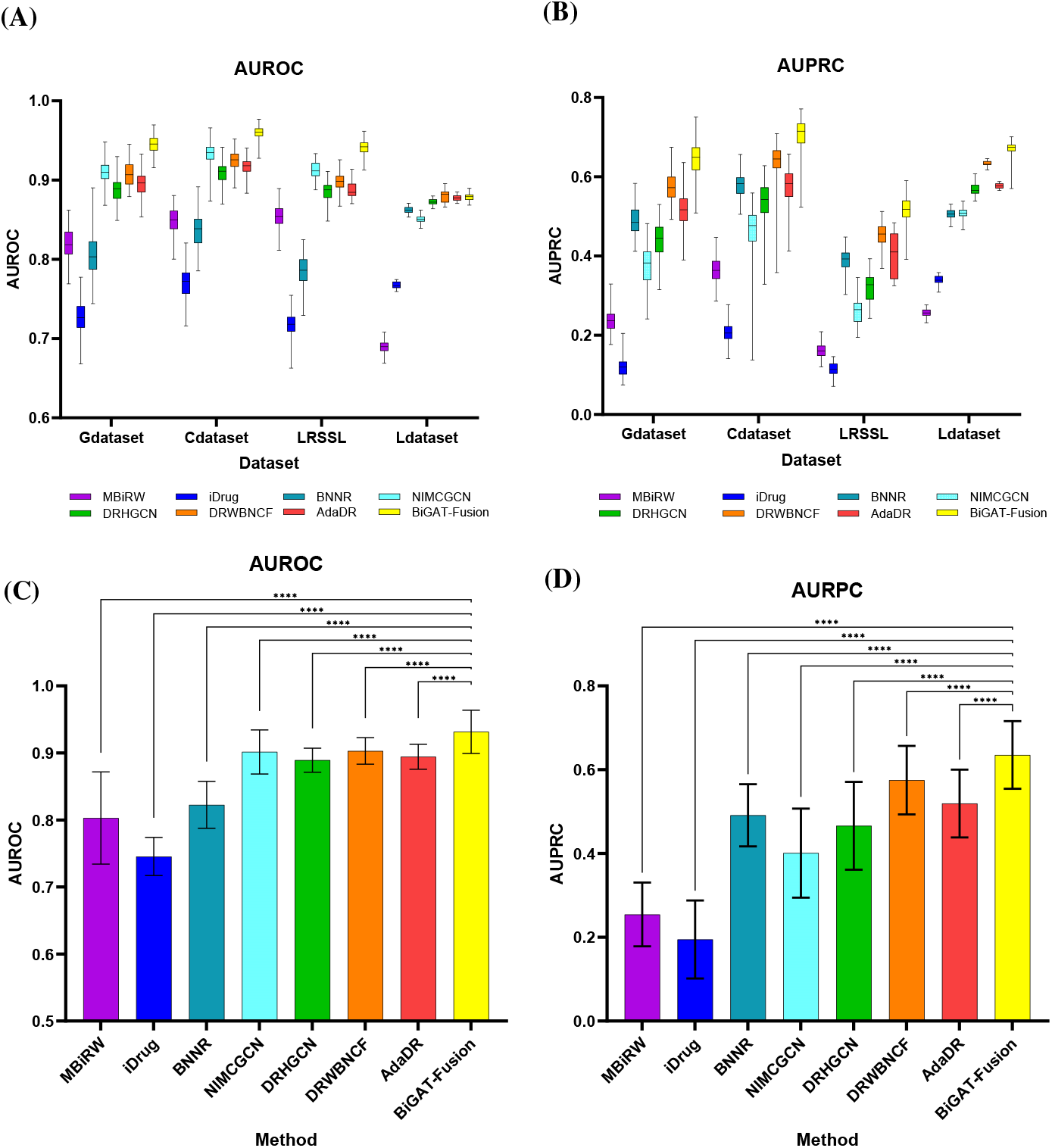
Visual comparison of model performance. **(A)** AUROC box–and–whisker plots across the four datasets (all runs per method). **(B)** AUPRC box–and–whisker plots across the four datasets (all runs per method). **(C)** AUROC summarized per method as bars; significance brackets indicate pairwise post-hoc comparisons against BiGAT-Fusion using one-way ANOVA with Bonferroni correction; **\*\*\*\*** denotes Bonferroni-adjusted *p* < 0.0001. **(D)** AUPRC summarized per method as bars; significance as in (C), one-way ANOVA with Bonferroni correction and **\*\*\*\*** for adjusted *p* < 0.0001.

Across all four datasets, BiGAT-Fusion attains the highest AUPRC. The mean scores are 0.647 ± 0.045 on Gdataset, 0.705 ± 0.045 on Cdataset, 0.516 ± 0.035 on LRSSL, and 0.673 ± 0.015 on Ldataset. The margins over the second-best method (DR-WBNCF) are +0.071, +0.066, +0.063, and +0.041, respectively. Given the severe class imbalance, these gains indicate a more reliable prioritization of true associations among many unknown pairs. BiGAT-Fusion also performs strongly on AU-ROC. It ranks first on Gdataset (0.945 ± 0.011), Cdataset (0.960 ± 0.008), and LRSSL (0.942 ± 0.008). On Ldataset, DRWBNCF achieves the highest AUROC (0.880 ± 0.008), while BiGAT-Fusion is a close second (0.879 ± 0.004), indicating comparable discrimination on this dataset.

Fig. 2(A)–(B) show box–and–whisker plots over all runs. The distributions for BiGAT-Fusion lie above those of the baselines on both metrics, with visibly tighter dispersion on several datasets. Statistical testing in Fig. 2(C)–(D) is reported at the method level across datasets: each box depicts, for a given method, the distribution of its scores over the four benchmarks (Gdataset, Cdataset, LRSSL, Ldataset). Significance brackets then compare each baseline against BiGAT-Fusion using one-way ANOVA with Bonferroni post-hoc comparisons; the anno-tation “**\*\*\*\***” denotes Bonferroni-adjusted *p* < 0.0001. Under this analysis, *all* baselines are significantly inferior to BiGAT-Fusion on both AUROC and AUPRC, indicating that our improvements hold consistently across datasets.

Overall, BiGAT-Fusion delivers state-of-the-art AUPRC together with competitive AUROC on all benchmarks. This behaviour is consistent with the model design: direction-aware bipartite attention captures asymmetric drug–disease signals, and node-wise gated fusion adapts the contribution of feature and topology views per node. Ablation evidence in Section 3.3 will further supports these choices.

### 3.3. Ablation Studies

We assess the contribution of each component using five configurations on all four datasets (Table 4): **BiGAT-Fusion (Full)**; **Scalar gate**, which replaces node-wise fusion with a single learnable scalar for drugs and for diseases; **Topology (no attention)**, which keeps the bipartite topology but substitutes directional attention with uniform mean aggregation; **Feature-graph only**, which removes the topology view and uses only the *k*NN feature graphs; and **Topology only**, which removes the feature graphs and uses only the bipartite topology.

**Table 4:** Ablation study of BiGAT-Fusion on four benchmark datasets.

| Model Variant | Gdataset |  |  | Cdataset |  |  | LRSSL |  |  | Ldataset |  |  |
| --- | --- | --- | --- | --- | --- | --- | --- | --- | --- | --- | --- | --- |
|  | AUROC | AUPRC | ΔAUPRC (%) | AUROC | AUPRC | ΔAUPRC (%) | AUROC | AUPRC | ΔAUPRC (%) | AUROC | AUPRC | ΔAUPRC (%) |
| **BiGAT-Fusion (Full)** | **0.945** | **0.647** | – | **0.960** | **0.705** | – | **0.942** | **0.516** | – | **0.879** | **0.673** | – |
| Scalar gate | 0.937 | 0.630 | ↓2.7% | 0.960 | 0.677 | ↓4.0% | 0.942 | 0.515 | ↓0.2% | 0.879 | 0.661 | ↓1.7% |
| Topology (no attention) | 0.910 | 0.483 | ↓25.4% | 0.934 | 0.546 | ↓22.5% | 0.898 | 0.377 | ↓27.0% | 0.862 | 0.606 | ↓9.9% |
| Feature-graph only | 0.912 | 0.487 | ↓24.7% | 0.936 | 0.562 | ↓20.2% | 0.920 | 0.366 | ↓29.2% | 0.854 | 0.536 | ↓20.2% |
| Topology only | 0.935 | 0.530 | ↓18.1% | 0.949 | 0.617 | ↓12.4% | 0.917 | 0.415 | ↓19.5% | 0.870 | 0.591 | ↓12.2% |

Values are means over the repeated 10-fold CV runs (rounded to three decimals). ΔAUPRC is the relative drop with respect to the Full model on the same dataset: Δ% = (AUPRC<sub>Full</sub> – AUPRC<sub>variant</sub>)/AUPRC<sub>Full</sub> × 100.

Removing directional attention or discarding one view yields the largest AUPRC losses. Relative to the full model, *Topology (no attention)* drops by 25.4% on Gdataset, 22.5% on Cdataset, and 27.0% on LRSSL; on Ldataset the decrease is 9.9%. *Feature-graph only* shows comparable declines of 24.7%, 20.2%, and 29.2% on Gdataset, Cdataset, and LRSSL, and 20.2% on Ldataset. These results indicate that direction-aware bipartite message passing is critical, especially under the extreme sparsity of LRSSL, and that attention remains beneficial on the denser Ldataset. Both views are necessary and complementary. *Topology only* underperforms the full model by 18.1% (Gdataset), 12.4% (Cdataset), 19.5% (LRSSL), and 12.2% (Ldataset) in AUPRC. Together with the “feature-only” variant, this pattern shows that chemical/semantic similarity and observed associations contribute distinct signals: topology dominates in sparse regimes, whereas similarity remains consistently valuable across datasets. Replacing node-wise fusion with *Scalar gate* produces smaller but consistent degradations. The AUPRC margins relative to the full model are 2.7% on Gdataset, 4.0% on Cdataset, 0.25% on LRSSL, and 1.7% on Ldataset. This confirms that per-node gating, which adapts the feature–topology balance at the node level, provides steady gains over a fixed, type-level blend.

In summary, the ablations support two design principles of BiGAT-Fusion: (i) direction-aware bipartite attention is essential for capturing the asymmetric drug → disease and disease → drug semantics; and (ii) adaptive, node-wise fusion reliably improves ranking by letting the model emphasize the right view for each node. These trends hold on AUROC as well and are consistent across datasets.

### 3.4. Interpretability Analysis

Having established through the ablation studies that direction-aware bipartite attention and node-wise gated fusion are the principal drivers of our gains, we now examine *why* these components help by analyzing the model’s internal signals on the training graph. All quantities are extracted exclusively on the training graph of each CV fold from the best checkpoint. For each dataset and fold we export: node-wise gates for drugs and diseases (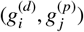) and directional attention weights on the bi-partite training graph, with the convention that 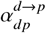 is the attention on edge *d* → *p* in the drug → disease direction and 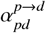 is the attention on *p* → *d* in the disease → drug direction. The table aggregates these fold-level artifacts as follows.

Results are reported as mean ± std. dev. The best result in each column is in **bold**, and the second-best is underlined.

Values are means over the repeated 10-fold CV runs (rounded to three decimals). ∆AUPRC is the relative drop with respect to the Full model on the same dataset: ∆% = (AUPRC_Full_ − AUPRC_variant_)/AUPRC_Full_ × 100.

- **Gate statistics**. We concatenate gate values across all folds and nodes of the same type and report mean ± std for drugs (*g*_*d*_) and diseases (*g*_*p*_).
- **Gate vs. degree (Spearman)**. Degrees are the *in-degrees on the training bipartite graph* (by direction). For drugs we use 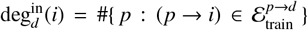 for diseases 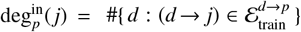. We pool nodes over folds and report Spearman’s ρ and its two-sided *p*-value for (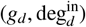) and (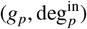).
- **Directional attention entropy**. Although attentions are normalized *per destination*, we summarize dispersion *per source node* in each direction: for drug→disease,

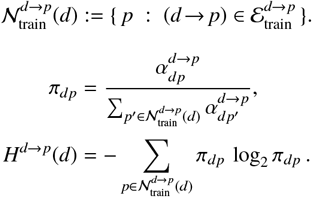

Analogously define *H*^*p*→*d*^(*p*) for the disease → drug direction. Degree-one sources have *H* = 0. For a paired, within-edge comparison we randomly sample up to *n*_sample_=1000 *train-ing* edges per fold, form pairs (*H*^*d*→*p*^(*d*), *H*^*p*→*d*^(*p*)) along the same edge (*d, p*), pool all pairs across folds, and report the mean entropies (as *H*_*d*→*p*_ and *H*_*p*→*d*_ in the table) together with a two-sided Wilcoxon signed-rank *p*-value.

Table 5 reports three complementary diagnostics computed on the training graph only: node-wise fusion gates, their correlation with node degrees, and the entropy of directional attentions. Together, these statistics clarify how BiGAT-Fusion balances feature and topology information and why directional message passing matters.

**Table 5:** Statistical summary of model interpretability mechanisms.

| Dataset | Node-wise Gate Statistics ($g_i$) | | | | Gate vs. Degree Correlation | | | | Directional Attention Entropy ($H$) | | |
| --- | --- | --- | --- | --- | --- | --- | --- | --- | --- | --- | --- |
| | Drug ($g_d$) | | Disease ($g_p$) | | Drug ($\rho_d, p_d$) | | Disease ($\rho_p, p_p$) | | $H_{d \rightarrow p}$ | $H_{p \rightarrow d}$ | p-value (Wilcoxon) |
| | Mean | Std. | Mean | Std. | $\rho$ | $p$ | $\rho$ | $p$ | Mean | Mean | |
| Gdataset | 0.476 | 0.061 | 0.421 | 0.042 | -0.157 | < 0.001 | -0.431 | < 0.001 | 1.594 | 3.001 | < 0.001 |
| Cdataset | 0.461 | 0.064 | 0.433 | 0.048 | -0.203 | < 0.001 | -0.448 | < 0.001 | 1.872 | 3.085 | < 0.001 |
| LRSSL | 0.475 | 0.032 | 0.450 | 0.036 | -0.294 | < 0.001 | -0.391 | < 0.001 | 1.882 | 2.685 | < 0.001 |
| Ldataset | 0.232 | 0.006 | 0.388 | 0.014 | 0.131 | < 0.001 | -0.198 | < 0.001 | 5.794 | 4.521 | < 0.001 |

Gate statistics ( $g_i$ ) and direction-wise attention entropies are computed *exclusively on the training graph* of each fold (constructed from positive edges in the training folds only; validation/test edges are excluded). $H_{d \rightarrow p}$ (**Drug**→**Disease**) and $H_{p \rightarrow d}$ (**Disease**→**Drug**) are Shannon entropies in base-2 (bits). Spearman’s $\rho$ measures correlation between gate values and node degrees (on the training graph). The Wilcoxon signed-rank $p$ -value compares per-edge entropies $\{H_{d \rightarrow p}, H_{p \rightarrow d}\}$ for the same sampled training edges. For readability, all $p$ -values smaller than 0.001 are reported as “< 0.001”.

We first examine the directional attention distributions. On **Gdataset, Cdataset**, and **LRSSL**, the mean entropy is consistently lower for Drug→Disease than for Disease→Drug (*H*_*d*→*p*_ < *H*_*p*→*d*_; Gdataset: 1.594 vs. 3.001; Cdataset: 1.872 vs. 3.085; LRSSL: 1.882 vs. 2.685; Wilcoxon *p* < 0.001 in all cases). This pattern indicates selective aggregation when updating diseases (few high-weight drugs) but broader pooling when updating drugs (many contributing diseases). On **Ldataset**, the asymmetry reverses (*H*_*d*→*p*_=5.794 vs. *H*_*p*→*d*_=4.521, *p* < 0.001), suggesting that the model adapts its directional selectivity to the denser association regime. These findings support the need for a direction-aware attention mechanism rather than uniform neighborhood averaging. We next consider the *node-wise fusion gates*. The gate value *g* controls the blend between feature-view and topology-view embeddings. Across Gdataset, Cdataset, and LRSSL the mean gates sit near 0.45–0.48 for both types, indicating a mild but consistent tilt toward topology on sparse graphs (e.g., Cdataset: *g*_*d*_=0.461 ± 0.064, *g*_*p*_=0.433 ± 0.048). On Ldataset, the drug gate is notably smaller (*g*_*d*_=0.232 ± 0.006) while the disease gate remains below 0.5 (*g*_*p*_=0.388 ± 0.014), reflecting a stronger reliance on topology where associations are more plentiful. Finally, we assess whether gating merely reflects node degree. Spearman correlations are small-to-moderate in magnitude (e.g., Cdataset: ρ_*d*_ = −0.203, ρ_*p*_ = −0.448; LRSSL: ρ_*d*_ = −0.294, ρ_*p*_ = −0.391; all *p* < 0.001), and mixed in sign on Ldataset (ρ_*d*_ = 0.131, ρ_*p*_ = −0.198, both *p* < 0.001). Thus, the gates are not a trivial function of connectivity. Instead, they encode content-driven preferences that adjust how much each node should trust topology versus features.

In summary, the entropy asymmetries confirm that the model learns *direction-specific* attention patterns, and the gating analysis shows *node-specific* fusion that is not reducible to degree effects. These mechanistic observations align with the ablation outcomes: removing attention and replacing it with uniform means erases the directional selectivity captured in the entropies and leads to the marked performance drops observed in Table 4.

Table 5 established three aggregate facts on **Gdataset**: lower directional dispersion for drug→disease than disease→drug (*H*_*d*→*p*_<*H*_*p*→*d*_; Wilcoxon *p*<0.001), gate means below 0.5 (a mild tilt toward topology), and non-trivial gate– degree correlations. Figure 3 complements these summaries with distributional, per-edge and per-node views that make the mechanisms visually explicit. Qualitatively similar phenomena hold on the remaining datasets; see Supplementary Fig. S1 (Cdataset), Fig. S2 (LRSSL), and Fig. S3 (Ldataset).

**Figure 3.**
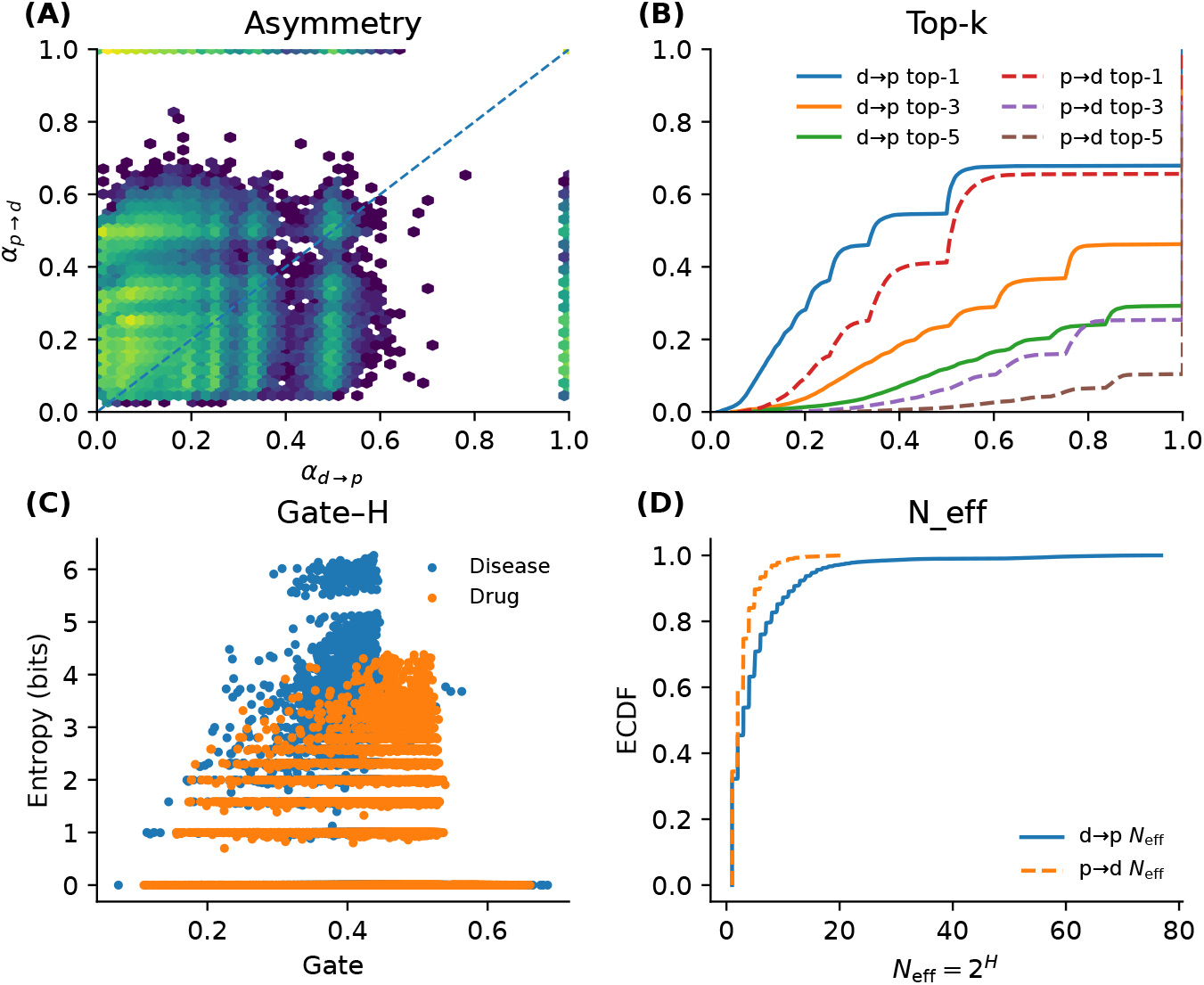
Interpretability on Gdataset. **(A) *Asymmetry***: per-edge bidirectional attentions; most points lie above the diagonal, indicating larger weights when aggregating to diseases (drug → disease) than to drugs (disease → drug). **(B) *Top-****k*: empirical cumulative distribution function (ECDF) of the cumulative incoming mass captured by the top-*k* neighbors per node; drug → disease curves are steeper, showing stronger concentration on few neighbors. **(C) *Gate–****H*: node-wise fusion gate versus attention entropy; gates vary smoothly across nodes, consistent with content-driven (not uniform) fusion. **(D)** *N*_eff_ : ECDF of the effective number of influential neighbors *N*_eff_ = 2^*H*^; both directions confirm that only a small subset of neighbors dominates.

All quantities are computed exclusively on the training graph of each cross-validation fold and then pooled across folds. Let 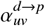 denote the attention weight on edge *u* → *v* when the desti-nation is a disease (drug→disease direction), and 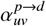 when the destination is a drug (disease → drug direction). In both views, the BiGAT layer applies a stabilized per-destination softmax, so that for any destination node *v*,

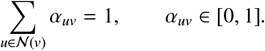

#### (A) Asymmetry

For each training edge (*d, p*) we pair the two directional weights on the same undirected pair: 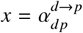 and 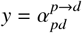, and plot the 2D density (hexbin) of (*x, y*).

#### (B) Top-*k* concentration

For each destination node *v*, collect its incoming weights {α_*uv*_}_*u*∈ 𝒩 (*v*)_, sort them in descending order *w*_(1)_(*v*) ≥ *w*_(2)_(*v*) ≥ ⃛, and define the cumulative top-*k* mass

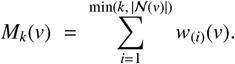

We then form the empirical cumulative distribution function (ECDF) 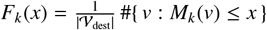 separately for drug→disease and disease→drug directions (we use *k* ∈ {1, 3, 5}).

#### (C) Gate–*H* coupling

For each node *v*, we compute the Shannon entropy (in bits) of its incoming attention distribution,

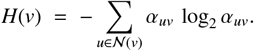

We pair *H*(*v*) with the learned node-wise fusion gate *g*_*v*_ (drug nodes use 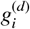, diseases 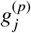; cf. Section 2.3) and display the scatter to examine how fusion adapts to dispersion.

#### (D) Effective number of neighbors

We summarize dispersion as an effective neighborhood size

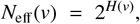

and plot its ECDF by direction. N_eff_ equals the number of equally weighted neighbors that would yield the same entropy, so larger values indicate more diffuse aggregation.

These construction details connect the distribution-level statistics in Table 5 to their concrete graphical origins in Fig. 3. The hexbin of paired directional weights (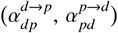) concentrates above the identity line, indicating that, on the same drug– disease pair, aggregation toward diseases typically assigns larger mass than aggregation toward drugs.

The hexbin of paired directional weights (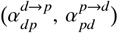) concentrates above the identity line, indicating that, on the same drug–disease pair, aggregation toward diseases typically assigns larger mass than aggregation toward drugs. The thin vertical and horizontal ridges at *x*=1 and *y*=1 arise from degree-one destinations, where the stabilized per-destination softmax assigns all probability to the solitary incoming edge. This edge-level asymmetry is the local counterpart of the entropy gap in Table 5. The ECDF of the cumulative top-k mass *M*_*k*_(*v*) shows markedly steeper curves for drug→disease than disease→drug. Hence, for many disease destinations a handful of neighbors (often the top-1 to top-3) already captures most of the incoming mass, whereas the mass entering drug nodes is spread more gradually across their neighbors. This pattern explains why replacing attention with uniform means hurts AUPRC on **Gdataset** (Table 4, “Topology (no attention)”): the model needs to upweight only a few highly informative neighbors. Plotting the learned node-wise gates *gv* against dispersion *H*(*v*) reveals a smooth, content-driven adaptation: when incoming topology is diffuse (larger *H*), gates tend to increase (larger *gv*), shifting weight toward the feature view; when topology is concentrated (smaller H), gates move down (smaller *gv*), relying more on the topology view. This behavior is consistent with the idea behind node-wise fusion and with the type-level means *g* ^*(d)*^ */g* ^*(p)*^ reported in Table 5. Importantly, the broad spread of points confirms that gating is not a trivial function of connectivity. The ECDF of N_eff_ (*v*) = 2^*H*(*v*)^ rises sharply at small values in both directions, indicating that most nodes effectively aggregate from only a few influential neighbors (small N_eff_), with a long but thin tail. This sparsity of effective support clarifies why directional attention improves ranking under class imbalance, and echoes the AUPRC gains over uniform aggregation in Table 4.

Overall, panels (A)–(D) explain the table-level findings mechanistically: asymmetric edge weights create lower *H* on the drug → disease direction, top-*k* curves and small *N*_eff_ show that only a small set of neighbors dominates, and node-wise gates adaptively shift between views depending on dispersion.

### 3.5. Hyper-parameter Sensitivity

We assess the robustness of BiGAT-Fusion to two operationally important hyper-parameters feature *k*-NN size (*k*) and in-batch negatives per positive (*k*_neg_). All other settings are fixed to the defaults in Section §2.4. Figure 4 summarizes mean: panels (A,B) report AUROC and (C,D) report AUPRC as each hyper-parameter is varied, with a single legend shown at the bottom.

**Figure 4.**
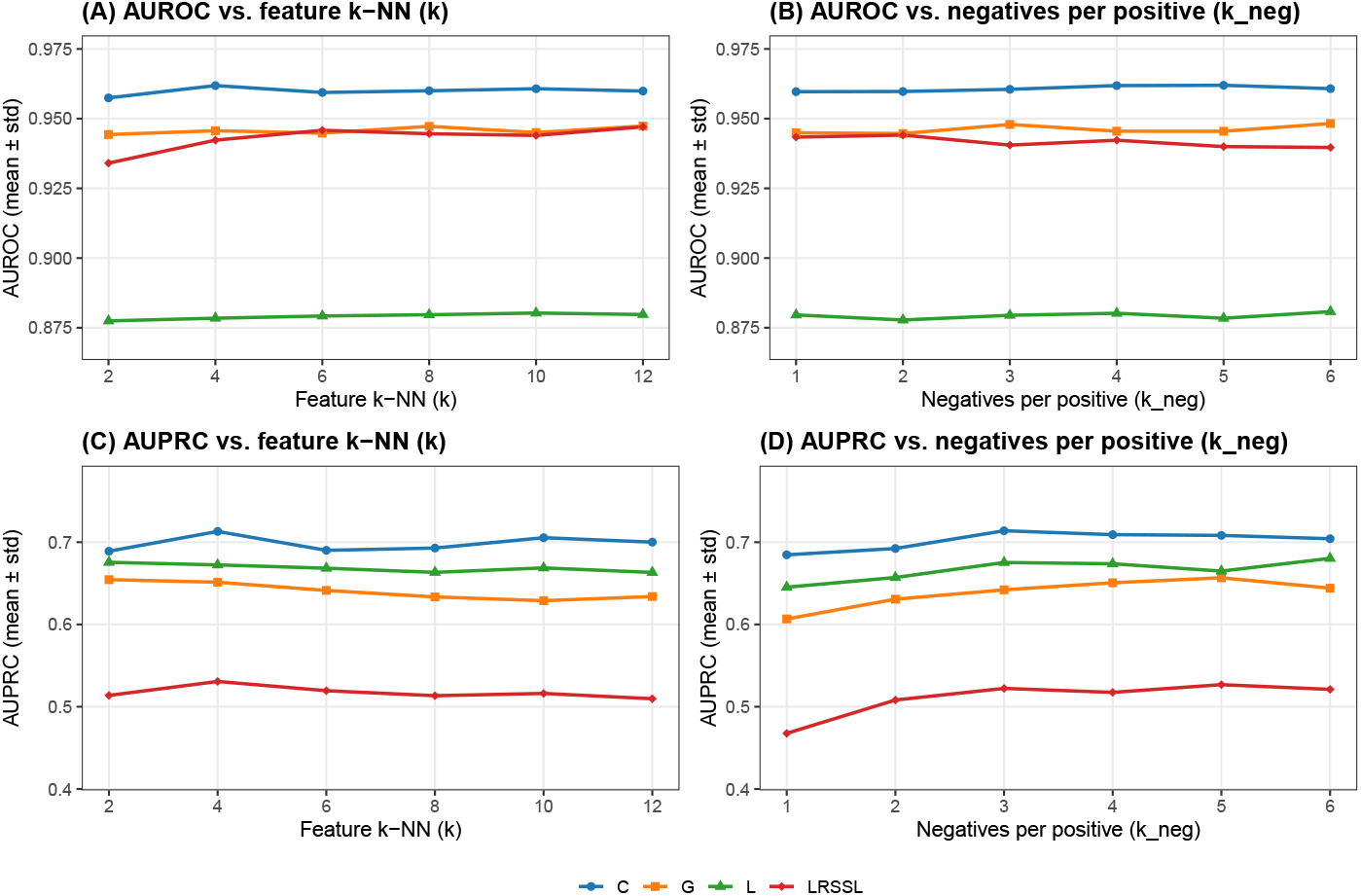
Hyper-parameter sensitivity. (A,B) AUROC vs. feature *k*-NN and *k*_neg_; (C,D) AUPRC vs. the same.

Feature *k*-NN (panels A,C) display a broad performance plateau across moderate *k*. Extremely small *k* reduces both AUROC and AUPRC (under-smoothing: too few informative neighbors in the feature view), while very large *k* also degrades performance (over-smoothing: excessive averaging washes out discriminative variation). Negatives per positive *k*_neg_ (panels B,D) display that AUROC varies weakly with *k*_neg_, whereas AUPRC is more sensitive: increasing *k*_neg_ beyond a small value typically yields diminishing or negative returns. This is consistent with the fact that *k*_neg_ sets the within-batch class prior for the BCE loss; a harsher prior (large *k*_neg_) can inflate apparent separability on ROC while hurting precision–recall in the highly skewed full-ranking task. In practice, small values (e.g., *k*_neg_ ∈ {3}) strike a good balance between gradient signal and PR-oriented ranking.

Collectively, (i) BiGAT-Fusion is stable to moderate perturbations of *k* and *k*_neg_, with AUROC largely flat and AUPRC forming a gentle peak around the defaults; (ii) extremes of either hyper-parameter are undesirable (under/over-smoothing for *k*, overly harsh batch prior for *k*_neg_); (iii) we therefore adopt *k*=4 and *k*_neg_=3 as dataset-agnostic defaults used throughout the paper.

### 3.6. Case Studies

To assess whether BiGAT-Fusion can surface clinically plausible hypotheses, we performed two case studies on Breast cancer (BC; OMIM:114480) and Small-cell cancer of lung (SCLC; OMIM:182280). For this analysis, the model was trained on the full set of known associations in Gdataset to emulate a realworld discovery setting, and we ranked all unknown drug candidates for each disease. The Top-10 lists were then cross-checked against guidelines, trial databases, and peer-reviewed literature (Table 6). The results are summarised below by evidence tier rather than implying current standard-of-care.

**Table 6:** External evidence for the Top-10 candidate drugs predicted for Breast cancer and Small-cell cancer of lung.

| Rank | DrugBank | Generic name | Key refs |
| --- | --- | --- | --- |
| *Case Study 1: Breast cancer (OMIM:114480)* |  |  |  |
| 1 | DB00305 | Mitomycin C | [26] |
| 2 | DB00399 | Zoledronic acid | [27] |
| 3 | DB00603 | Medroxyprogesterone acetate | [28, 29] |
| 4 | DB01196 | Estramustine phosphate | [30, 31] |
| 5 | DB00977 | Ethinyl estradiol | [32, 33] |
| 6 | DB01128 | Bicalutamide | [34, 35] |
| 7 | DB00541 | Vincristine | [36] |
| 8 | DB00290 | Bleomycin | [37] |
| 9 | DB00642 | Pemetrexed | [38] |
| 10 | DB01005 | Hydroxyurea | [39] |
| *Case Study 2: Small-cell cancer of lung (OMIM:182280)* |  |  |  |
| 1 | DB00290 | Bleomycin | [40] |
| 2 | DB01181 | Ifosfamide | [41] |
| 3 | DB00997 | Doxorubicin | [42] |
| 4 | DB00262 | Carmustine | [43] |
| 5 | DB00541 | Vincristine | [44] |
| 6 | DB00958 | Carboplatin | [45] |
| 7 | DB00851 | Dacarbazine | [46] |
| 8 | DB01073 | Fludarabine | [47] |
| 9 | DB00707 | Porfimer sodium | [48, 49] |
| 10 | DB00570 | Vinblastine | [50, 51] |

Key references point to clinical guidelines, reviews, trials, or other authoritative sources supporting the drug’s relevance to the disease.

Among the Top-10, several agents have guideline- or meta-analysis–level support in specific adjuvant settings (e.g., zoledronic acid), while others are cytotoxic components that have been historically investigated in multi-agent regimens (e.g., mitomycin C, vincristine) or evaluated in selected contexts (e.g., medroxyprogesterone acetate, estramustine). Thus, the list mixes agents with established adjuvant roles and drugs with historical or context-dependent evidence. For SCLC, the model highlights several classical cytotoxics. Notably, carboplatin—a core component of contemporary first-line regimens—appears among the top predictions. The list also recovers drugs that feature in older combination protocols (e.g., doxorubicin and vincristine in the CAV regimen), together with agents with limited or investigational evidence in SCLC.

Across both diseases, BiGAT-Fusion retrieves drugs with varying strength of external support, including guideline-backed agents (e.g., zoledronic acid for BC, carboplatin for SCLC) and historically studied or context-specific cytotoxics. This pattern is consistent with the model’s goal in a discovery setting: to prioritise plausible candidates that can be triaged by evidence instead of reproducing only the current standard-of-care.

Figure 5 visualises the Top-10 unknown candidates for each disease as a bipartite association network. Two patterns are evident. First, several predictions are *shared* across Breast cancer and Small-cell cancer of lung, most notably *vincristine* and *bleomycin*, and are connected to both diseases with relatively thick arcs, reflecting high scores. Second, each disease also exhibits *disease-specific* candidates with strong model support: for BC these include adjuvant-context agents such as *zoledronic acid* and *medroxyprogesterone acetate*, whereas for SCLC the model prioritises classical cytotoxics such as *carboplatin, doxorubicin*, and *carmustine*. This network view is consistent with Table 6 and our evidence-tier summary: BiGAT-Fusion retrieves both guideline-supported drugs (e.g., zoledronic acid for BC; carboplatin for SCLC) and historically investigated components (e.g., vincristine, bleomycin), thereby surfacing clinically plausible hypotheses that can be triaged by external evidence rather than merely reproducing the current standard-of-care.

**Figure 5.**
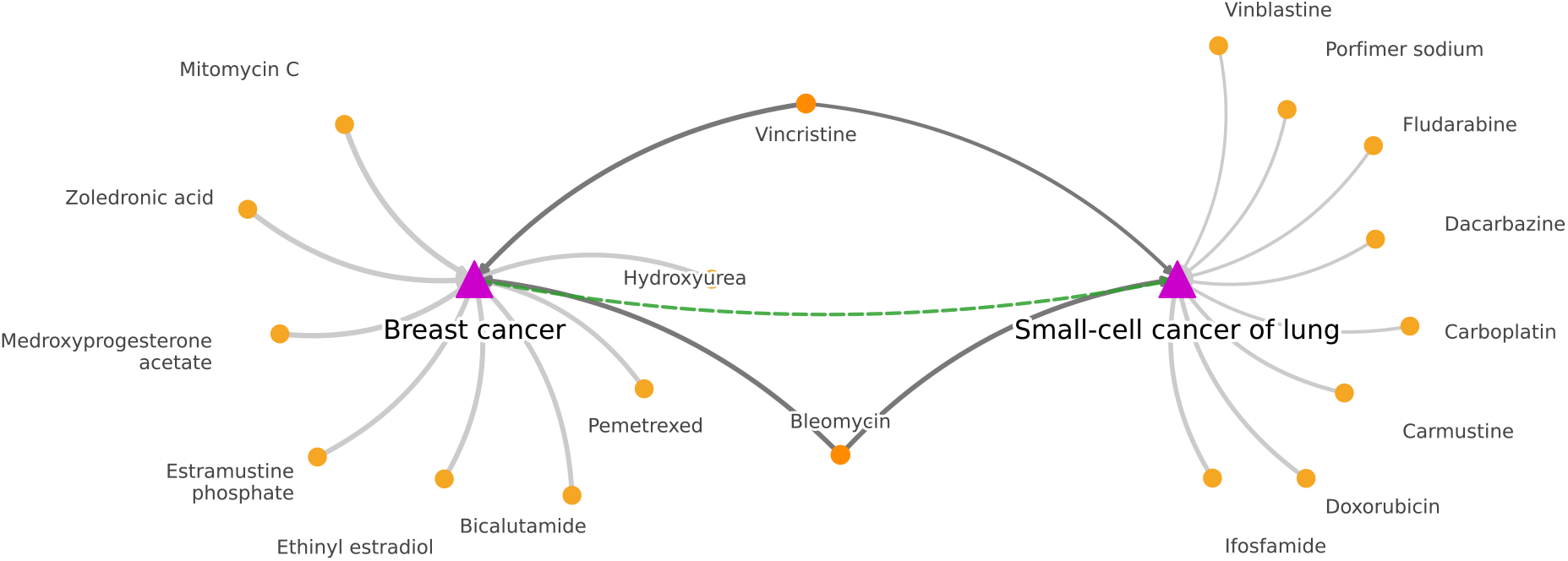
Top-10 candidate–drug association network for Breast cancer (left) and Small-cell cancer of lung (right). Magenta triangles denote diseases; orange circles denote drugs. Curved arrows point from drugs to diseases; thicker arcs indicate higher model-assigned probabilities. The green dashed bridge highlights the concept of “shared” top-ranked candidates predicted for both diseases.

## 4. Conclusion

In this work we introduced BiGAT-Fusion, a two-view graph model that combines direction-aware bipartite attention with type-specific, node-wise gated fusion to prioritize drug–disease associations under severe class imbalance. Across four widely used benchmarks, BiGAT-Fusion consistently delivered state-of-the-art AUPRC while remaining competitive in AUROC, reflecting improvements that matter for highly skewed discovery problems where precision–recall analysis is the more informative yardstick [52]. Mechanistically, analyses of directional attention entropy and per-node gates corroborated our design choices: the model learns asymmetric, destination-specific aggregation on the bipartite graph and adapts the contribution of feature versus topology evidence at the node level.

Methodologically, BiGAT-Fusion unifies strengths from several lines of DDA research. Early similarity- and network-based approaches (e.g., PREDICT and bi-random-walk formulations) showed that integrating drug–drug and disease–disease similarity with observed associations is a powerful bias for repurposing [8, 9]. Matrix-completion methods further exploited low-rank structure with principled regularization [10]. More recent GNNs generalized message passing to heterogeneous biomedical graphs, leveraging architectures such as GCNs and GATs [11, 12]. Our contribution complements this trajectory by (i) explicitly modeling directional asymmetry in bipartite aggregation, and (ii) learning node-wise fusion between feature-view signals defined on kNN graphs and topology-view signals induced by observed DDAs. The resulting Residual-MoE decoder improves pair scoring stability under class imbalance.

Practically, BiGAT-Fusion is designed to be a drop-in component for computational drug repurposing pipelines [3, 4]. The feature view relies on established similarity choices—such as extended-connectivity fingerprints with Tanimoto similarity for drugs and ontology/text-derived similarities for diseases—whose empirical utility is well supported [53, 54]. The topology view is constructed strictly from training positives to avoid structural leakage, and repeated K-fold cross-validation with validation-driven selection provides robust estimates of generalization.

This study also highlights limitations and opportunities. First, the quality of similarity inputs remains a bottleneck; richer, complementary modalities (targets, pathways, gene expression, clinical narratives) could be incorporated in the feature view to improve transfer to under-studied entities. Second, while our evaluation emphasizes pair ranking, prospective validation and task-specific calibration are essential for decision-support settings. Finally, beyond transductive setups, extensions to stricter cold-start regimes (held-out drugs/diseases), temporal splits, and uncertainty quantification would increase practical reliability. We anticipate that BiGAT-Fusion’s interpretable gating and directional attention mechanisms will remain valuable as larger knowledge graphs and multimodal evidence become standard in in silico repurposing.

## (Supplementary Material: S1)

**Figure S1:**
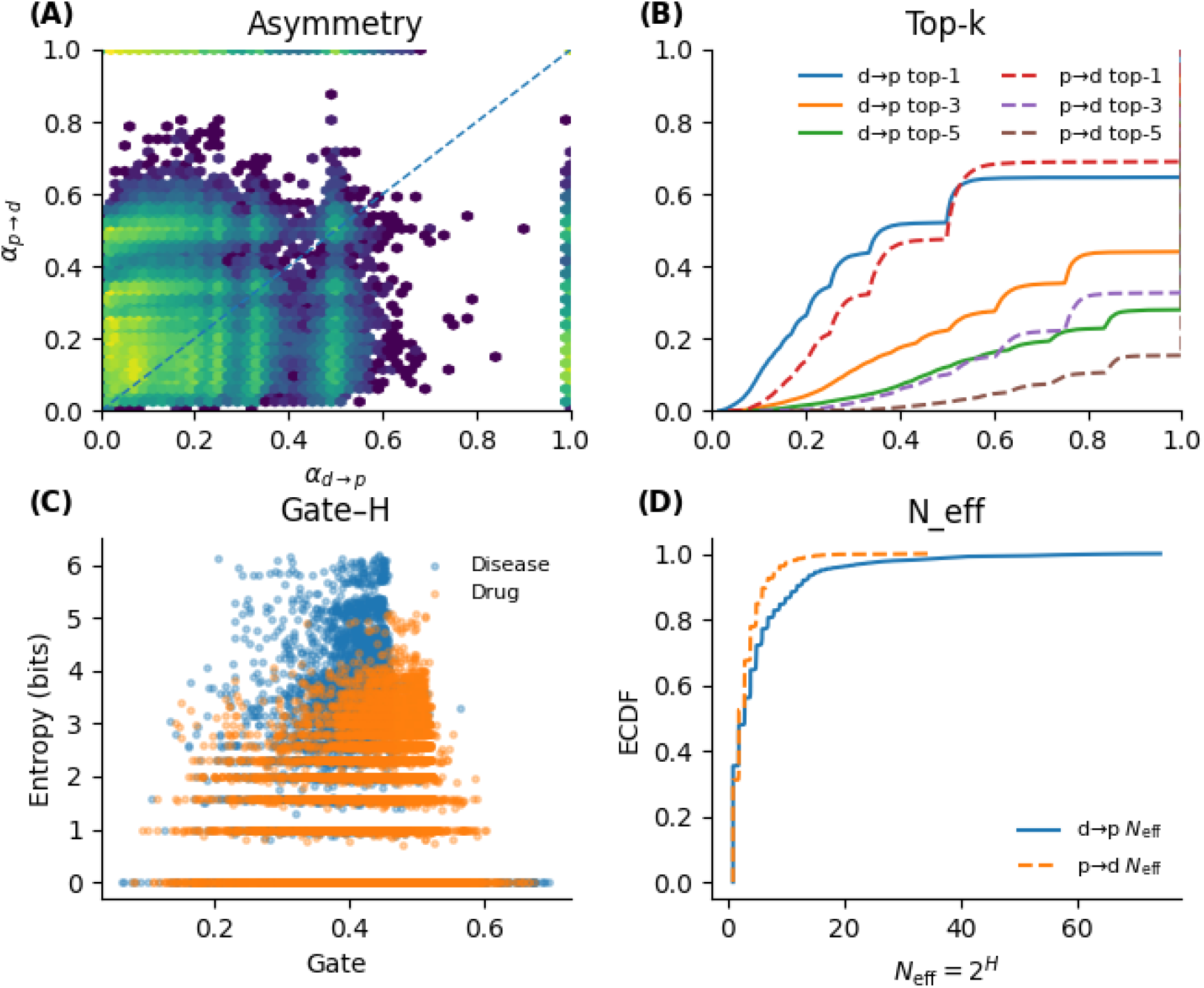
Supplementary interpretation results on Cdataset.

## (Supplementary Material: S2)

**Figure S2:**
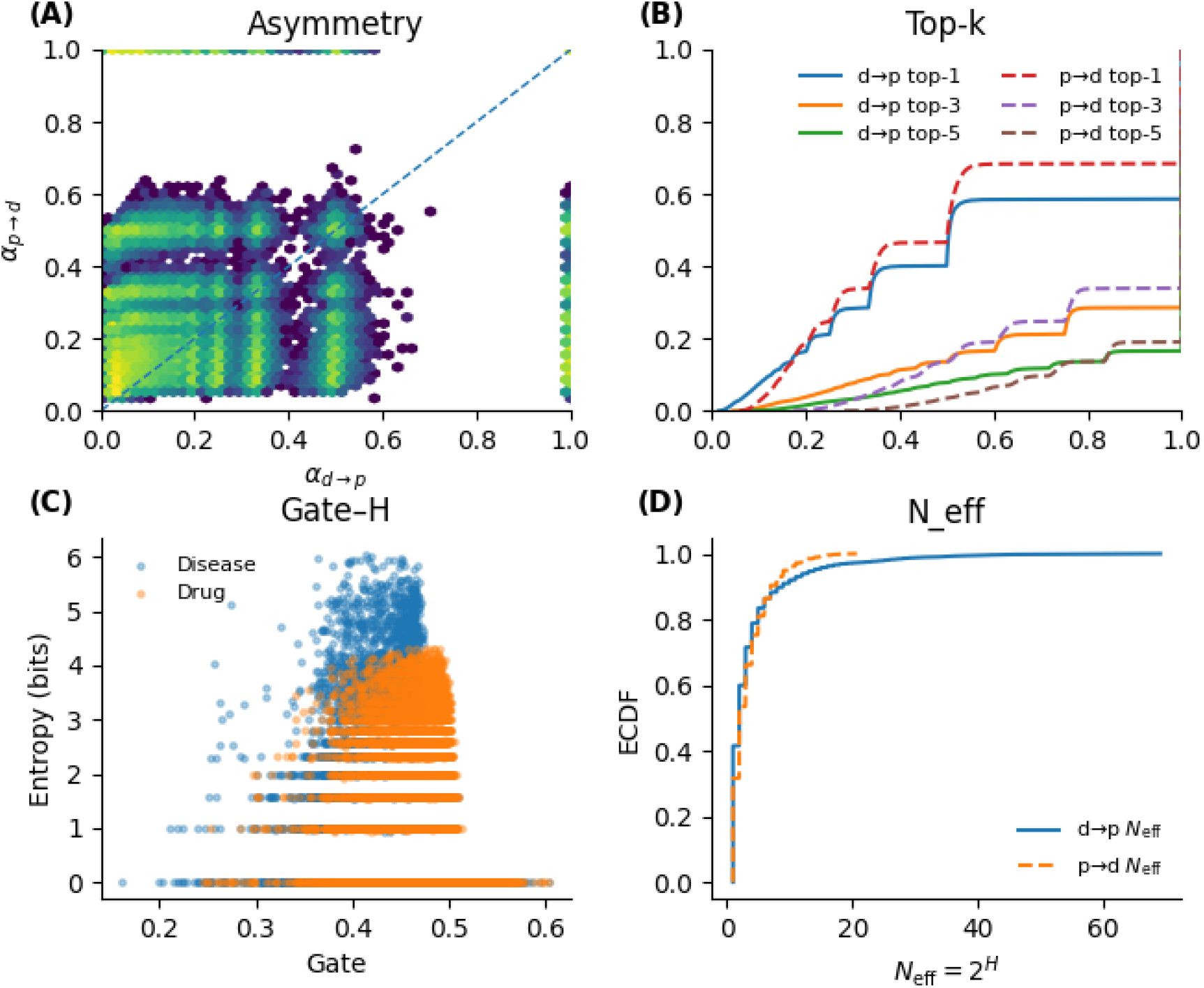
Supplementary interpretation results on LRSSL.

## (Supplementary Material: S3)

**Figure S3:**
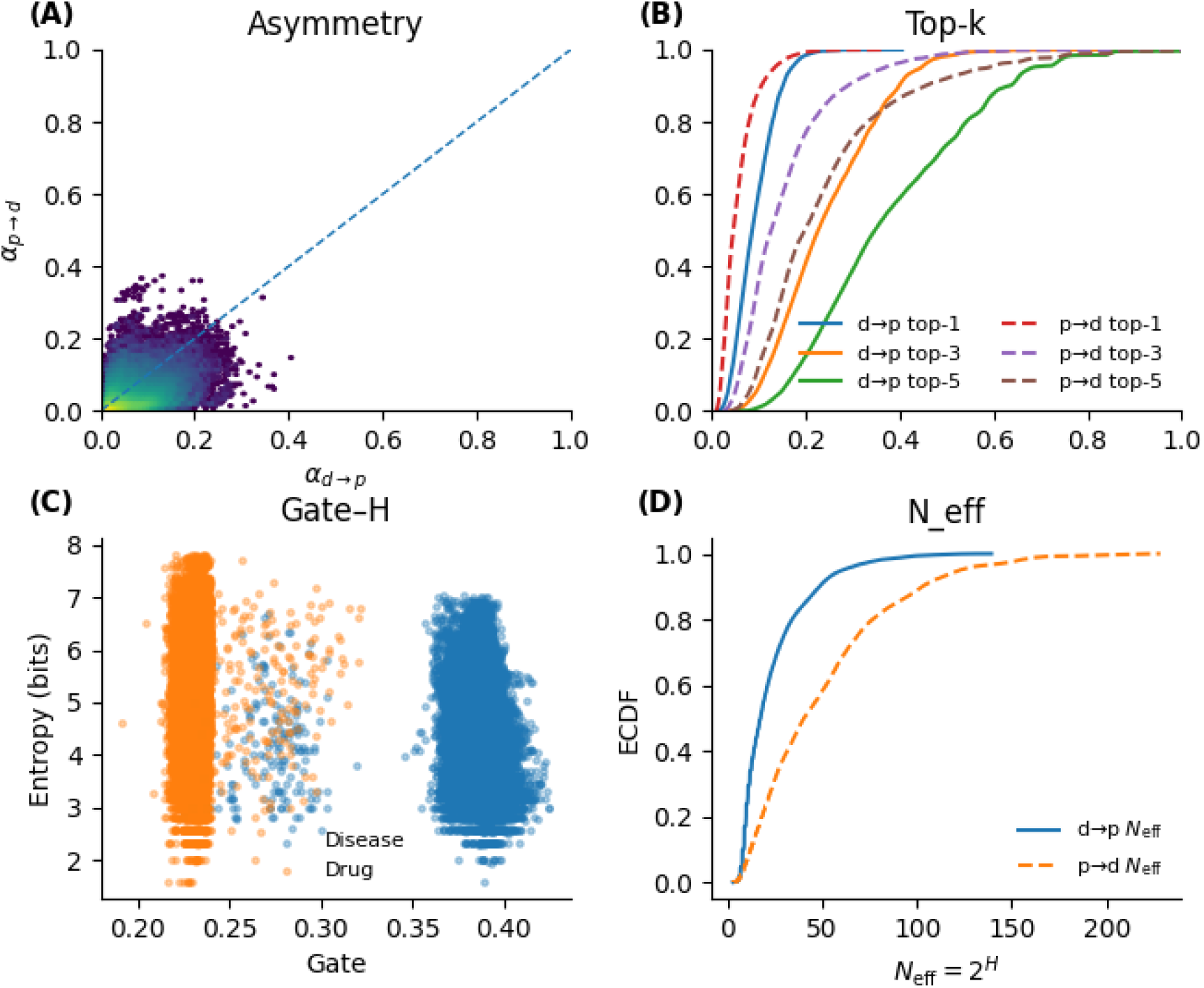
Supplementary interpretation results on Ldataset.

